# Neutrophil-neuronal crosstalk drives arthritis-induced pain

**DOI:** 10.64898/2025.12.27.696652

**Authors:** Zerina Kurtović, Juan Antonio Vazquez Mora, Sijing Ye, Sara Dochnal, Katalin Sandor, Mohd Sami Ur Rasheed, Sven David Arvidsson, Alex Bersellini Farinotti, Nilesh Agalave, Gustaf Wängberg, Matthew A. Hunt, Nils Simon, Julia Dorothea Monika Döring, Alexandra Kuliszkiewicz, Lizeth Ponce Gomez, Giovanni Emmanuel López Delgado, Arisai Martinez Martinez, Eduardo Mendoza Sanchez, Khosiyat Makhmudova, Enriqueta Munoz Islas, Miriam Bollmann, Mattias Svensson, Emerson Krock, Lisbet Haglund, Juan Miguel Jimenez Andrade, Tony L. Yaksh, Harald Lund, Camilla I. Svensson

## Abstract

Pain in rheumatoid arthritis (RA) often persists despite effective control of inflammation, suggesting distinct mechanisms driving nociception. In both patients and animal models, pain severity does not strongly correlate with the degree of inflammation^1,2^. Sensory neurons, with cell bodies located in the dorsal root ganglia (DRG), innervate peripheral tissues, including joints, and transmit pain signals to the central nervous system. Crosstalk between sensory neurons and immune cells occurs at all of these sites. While sensory neurons can be directly activated by immune mediators, it remains unclear whether pain-like behaviour in antibody-induced arthritis models arises independently of immune cell activity, or which immune cell populations and mediators are required to activate pronociceptive mechanisms. Through temporal profiling of the CAIA joint–DRG transcriptomic axis, we identified SEMA4D and OSM signalling as candidate molecular mediators of neutrophil–neuron communication and neuronal sprouting. The joint–DRG atlas also revealed persistent changes in the fibroblast-immune cellular composition of the joint, along with molecular changes in DRG neurons. We showed that mechanical and cold hypersensitivity, as well as sprouting of CGRP+ nociceptive fibers in synovial tissue of mice with collagen antibody-induced arthritis (CAIA), require neutrophils but not macrophages. Analysis of publicly available datasets showed that neutrophils from the synovium of RA patients express high levels of SEMA4D and OSM, and corresponding expression of their receptors, PLEXINB1 and OSMR, in human DRG neurons, underscoring the translational relevance of this axis. Both murine and human-derived DRG neurons sprout in response to OSM. Our findings demonstrate that neutrophils produce molecules that act as cues for nociceptor sensitization and structural remodelling. Targeting these molecules could improve the efficacy of RA treatments by reducing pain while simultaneously preventing disease progression.

## Introduction

Chronic joint pain is a defining feature of rheumatoid arthritis (RA), a prevalent autoimmune condition that affects approximately 0.5–1% of the global population ^3^. Although disease-modifying antirheumatic drugs (DMARDs) effectively reduce inflammation and alleviate pain, 40-50% of RA patients continue to report significant pain even after one year of treatment^1^. This discrepancy highlights the importance of identifying mechanisms driving RA-related pain and discovering novel therapeutic targets to simultaneously address inflammation and pain.

Antibody-induced arthritis models, such as the collagen antibody–induced arthritis (CAIA) model, replicate key features of rheumatoid arthritis–associated pain, including the persistence of pain-like behaviours beyond the resolution of joint inflammation^2,4^. This chronic hypersensitivity is associated with neuropathic-like changes in sensory neurons and sprouting of neuronal fibers within synovial tissues^5,6^, yet the signalling pathways that initiate and sustain these changes remain poorly defined.

Joint-innervating nociceptive neurons have their cell bodies in the dorsal root ganglia (DRG) adjacent to the spinal cord. They convey peripheral signals from the joint to the dorsal horn of the spinal cord, where this information is relayed to supraspinal centers for pain perception. Neuroimmune interactions occur at multiple levels along this axis, including within both the joint and the DRG^7–11^. Over the past years, single-cell transcriptomic profiling has provided detailed maps of the cellular composition of these tissues^12–14^. These studies have revealed that the DRG harbors a range of immune cell populations, including monocyte-derived macrophages, long-lived tissue-resident macrophages, neutrophils, and additional immune subsets. In parallel, sensory neurons have been subdivided into increasingly refined subtypes based on molecular identity, expanding beyond classical definitions based on size, myelination, and neurotransmitter usage^12,15–17^. The joint microenvironment has similarly been mapped in both healthy and arthritic states, revealing disease-associated shifts in macrophage and fibroblast populations^13,14,18–21^. However, a comprehensive, temporal atlas of the joint–DRG axis, specifically addressing neuroimmune communication during the development and persistence of arthritis-associated pain has yet to be established.

Macrophages are resident phagocytic immune cells with essential roles in both the induction and resolution of inflammation, adopting diverse functions based on their tissue niche^22,23^. Single cell analyses across multiple organs have identified three overarching subtypes of tissue-resident macrophages: embryonically derived macrophages expressing Timd4, Lyve1, or Folr2 (TLF); monocyte-derived macrophages expressing Ccr2; and MHCII^high^ macrophages (TLF- Ccr2-) of mixed origin^24^. Within the synovium, macrophage subsets are spatially organized into specific niches, such as lining and sublining macrophages^13^. This cellular heterogeneity in both mouse and human synovium is reflected in the diverse and sometimes opposing roles of synovial macrophages in arthritis, ranging from protective to pathogenic^13,19,20^. The role of these populations in the induction and maintenance of pain-like behaviour remains poorly understood. Selective depletion of CSF1R-expressing cells can alleviate mechanical hypersensitivity in an osteoarthritis model^9^ but prolong hypersensitivity in a monoarthritis model induced by Complete Freund’s Adjuvant^10^, highlighting the complexity of macrophage-mediated nociception.

Neutrophil infiltration into arthritic joints is well-documented in both patients and animal models^13,18,25^. Recent studies have also demonstrated the presence of neutrophils in DRG tissues^26^, although their functional role at this site remains understudied. Neutrophil infiltration into DRG has been observed in models of neuropathic pain and autoimmune encephalomyelitis (EAE)^27–29^. However, the functional consequences of neutrophil depletion appear to be context dependent. In EAE, depletion of neutrophils reverses mechanical hypersensitivity^28^, whereas following sciatic nerve injury^29^, increased infiltration of neutrophils has been associated with reduced thermal hypersensitivity. The exact contribution of neutrophils to neuronal remodelling and pain signal transmission in antibody-induced arthritis remains to be elucidated.

We hypothesized that immune cells initiate persistent pain behaviours and sensory neuron sprouting in antibody-induced arthritis. To investigate this, we generated a temporally resolved joint–DRG single-cell atlas capturing the progression from acute inflammation to resolution and the associated states of mechanical hypersensitivity. By integrating this atlas with targeted immune cell depletion experiments, we identified neutrophils as key drivers of joint remodelling during the inflammatory phase, with effects that persists beyond resolution. Notably, this atlas revealed neutrophil-derived factors such as SEMA4D and OSM as candidate factors promoting neuronal sprouting in the CAIA model. Importantly, these pathways are conserved in human RA, highlighting their potential as therapeutic targets for chronic inflammatory pain.

## Results

### Limited cellular remodelling in DRG during CAIA

To elucidate the neuroimmune mechanisms underlying the induction of mechanical hypersensitivity and its transition to chronicity, we profiled CAIA-induced changes in the joint-DRG axis using single cell transcriptomics. We performed 10X Genomics single-cell RNA-sequencing of DRG and joint tissues incorporating oligonucleotide-conjugated antibodies against MHCI and CD45 to label immune and non-immune compartments. Cells were isolated from L3–L5 DRGs and ankle joints from saline and CAIA mice at three time points capturing distinct disease phases, peak inflammation with mechanical hypersensitivity (day 10), resolution of inflammation with persistent mechanical hypersensitivity (day 20), and sustained mechanical hypersensitivity after resolution of overt inflammation (day 58) (Fig. 1A). Data was processed using the *Seurat* pipeline for demultiplexing and clustering, generating a temporal single-cell atlas of the joint–DRG axis in CAIA. Cell clusters were annotated based on established cell-type-specific marker gene expression.

**Figure 1.**
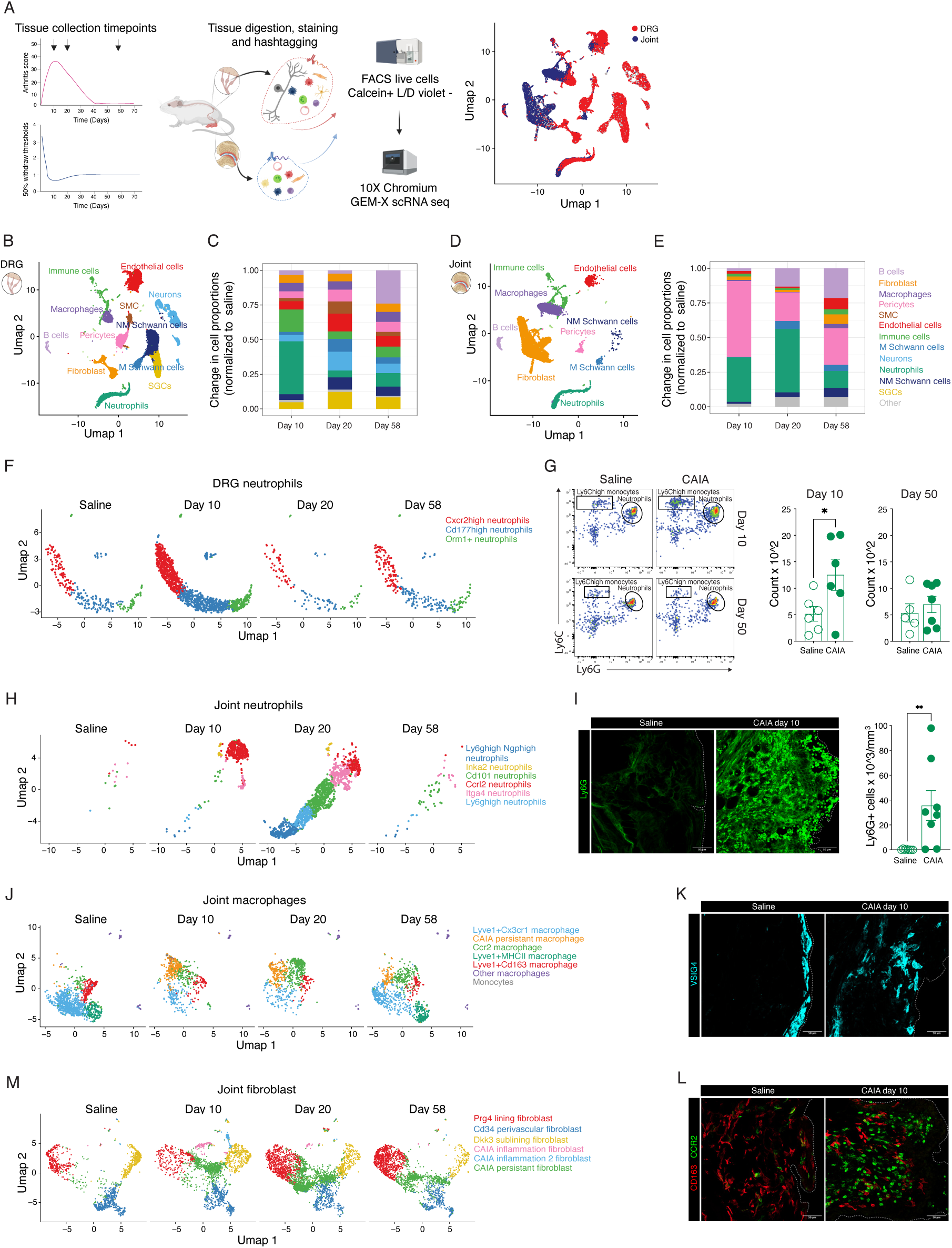
Molecular and cellular atlas of the joint-DRG axis in CAIA mice. (A) Design of sequencing experiment and Umap of sequenced cells form L3-L5 DRGs and hind paw joints of saline and CAIA mice at 3 timepoints. Four animals per group were pooled into 2-3 hashtagged samples. Created with BioRender. (B) Umap of cellular composition of the DRG in saline and CAIA. (C) Distribution of DRG cell types across timepoints normalized to saline. (D) Umap of the cellular composition of the joint in saline and CAIA mice. (E) Distribution of joint cell types across timepoints. (F) Reclustering of DRG neutrophils shows presence of three clusters over all timepoints: Cxcr2^high^ (red), Cd177^high^ (blue) and Orm1+ (green). (G) Flow cytometery analysis of DRG immune cells and quantification of Ly6G+ neutrophils on day 10 and day 58. (H) Reclustering of Joint neutrophils shows timepoint-dependent clusters: Ccrl2+ on day 10 (red), and Itga4+ (pink), Cd101+ (green), Ly6g^high^ (light blue) and Lyg6g^high^ Ngp^high^ (dark blue) on day 20 and the same subtypes were present at low numbers on day 58. (I) Stainings and quantification of neutrophils in the synovium of CAIA mice on day 10. (J) Reclustering of joint macrophages. (K) Stainings of VSIG4+ lining macrophages macrophage subtypes in the ankle joint synovium of saline and CAIA mice (day 10). (L) Stainings of CD163+ and CCR2+ sublining macrophages in the ankle joint synovium of saline and CAIA mice (day 10). (M) Reclustering of synovial fibroblasts. Unpaired t-test were used for comparisons. *P<0.05, **P<0.01

Beyond neurons, we detected all major cell types previously described in DRGs ^26,30,31^ including satellite glial cells (*Fabp7*), Schwann cells (*Ncmap*, *Scn7a*), macrophages (*Cd64*, *Cx3cr1*), endothelial cells (*Pecam*), pericytes (*Anpep*), fibroblasts (*Dcn*, *Pdgfra*), neutrophils (*Ly6g*), and other immune cells (*Ptprc*) (Fig. 1B, C, S1A). In contrast to the joint, where extensive cellular remodelling occurred subsequent to induction of CAIA (Fig. 1D, E, S1B), the overall cellular composition of the DRG remained largely stable (Fig. 1B, C). The only notable change was a transient expansion of DRG neutrophils at day 10 (Fig. 1C, F).

This increase spanned multiple neutrophil subtypes, which were also present in saline-injected mice (Fig. 1F). Reclustering of DRG neutrophils revealed three distinct clusters, two of which formed a temporal continuum characterized by decreasing *Cxcr2* and increasing *Cd177* expression (Fig. S1C), a pattern mirrored in joint-infiltrating neutrophils (Figure S1D). Additionally, we identified a subset of DRG neutrophils expressing *Orm1* (orosomucoid 1), *Cd55* (decay-accelerating factor), *Ptgr1* (prostaglandin reductase 1) and *Chil3* (chitinase-like protein 3), consistent with an immunomodulatory phenotype; notably, this population was already present in saline-injected mice (Fig. 1F, S1E). Immunostaining revealed that the majority of neutrophils resided in the spinal meninges (Fig. S1F, G), extending over the DRG parenchyma and into the epineurium at the distal end, in agreement with previous reports^32^. Although neutrophils were largely excluded from the DRG parenchyma, this compartment is exposed to mediators produced in the spinal meninges, and CSF perfuses the DRG parenchyma^32,33^. Notably, collagen II, the antigen target in CAIA, is present in the spinal meninges and colocalizes with neutrophils (Fig. S1G), potentially contributing to the observed neutrophil expansion in this region.

Despite these limited cellular changes, differential gene expression analysis revealed widespread transcriptional alterations across timepoints and cell types (Fig. S1H-I), with neurons showing the largest number of differentially regulated genes. To independently validate neutrophil infiltration, we performed flow cytometry on cervical and lumbar DRGs using an immune panel optimized for the DRG niche. This confirmed a transient increase in neutrophil (CD45+ CD11b+ Ly6G+) at day 10, with no sustained elevation at day 58 (Fig. 1G). A modest, transient increase in monocytes (CD45+ CD11b+ Ly6C+) infiltration was also detected at day 10 (Fig. S2A, S2B), while other DRG macrophage populations remained unchanged (Fig. S2B).

### CAIA leads to long-term alterations in the joint cellular niche

Next, we examined the joint cells in our dataset, which revealed a profound reorganization of joint architecture in CAIA (Fig. 1D, E). Compared to saline joints, CAIA joints exhibited a marked influx of neutrophils and monocytes, along with a reorganization of tissue-resident macrophages and fibroblasts. This included the emergence of CAIA-specific fibroblast clusters, and an increased number of pericytes (Fig. 1E). Notably, we also observed an unexpected influx of B cells (Fig. 1E), a surprising finding given that the CAIA model is generally considered independent of adaptive immune responses.

While neutrophils were virtually absent in the joints of saline-injected animals, neutrophils accumulating in CAIA joints displayed a shift in phenotype over time (Fig. 1H). *Ccrl2+ Cxcr2^high^* neutrophils predominated at day 10 and progressively transitioned into *Cd177+* subclusters expressing *Itga4, Cd101* or *Ngp* (Fig. 1H, S3A). *Cxcr2+* and *Cd177+* neutrophils have previously been identified in single-cell analysis of inflammatory lung and liver models. While CXCR2 is well established as a key regulator of neutrophil recruitment, CD177+ neutrophils exhibit enhanced neutrophil extracellular trap (NET) formation compared to their CD177- counterparts^34^. Using immunohistochemistry and confocal microscopy, we confirmed robust neutrophil infiltration of the CAIA ankle joint synovium at the peak of inflammation (day 10; Fig. 1I).

Reclustering of joint macrophages enabled annotation of distinct synovial macrophage populations previously described in the literature ^13^, including *Lyve1+ Cx3cr1+* lining macrophages, *Lyve1+ Cd163+* sublining macrophages, *Lyve1+ MHCII+* interstitial precursor macrophages, and monocyte-derived *Ccr2+* macrophages (Fig. 1J). During CAIA, we observed significant changes in the macrophage composition, with differential effects across synovial subsets. In particular, *Cd163+* sublining and *Cx3cr1+* lining macrophages were reduced, consistent with histological evidence of lining disruption, as indicated by the loss of the lining marker VSIG4, alongside a concomitant expansion of *Ccr2+* macrophages (Fig. 1J-L). Interestingly, a subset of *Ccr2+* macrophages formed a CAIA-specific cluster (Fig. 1J). Importantly, even at day 58, the macrophage landscape had not fully reverted to that seen in joints of saline injected mice, indicating that CAIA induces persistent alterations in synovial immune cell composition (Fig. 1J).

Similarly, subsetting and reclustering joint fibroblasts revealed distinct populations in agreement with previous studies^14,35^, including *Prg4+* lining fibroblasts, two *Thy1+* sublining fibroblast populations characterized by either *Cd34+* or *Dkk3+* expression, and three CAIA-specific clusters, two present during the inflammatory phase and one persisting into the late phase (Fig. 1M).

Differential gene expression analysis of joint cells across time points revealed extensive transcriptional changes. Mapping these differentially expressed genes back onto their respective cell types highlighted broad molecular alterations throughout the joint, with synovial fibroblasts exhibiting the most pronounced changes (Fig. S3A). Neutrophil-related genes (*Ly6g, Ngp, Cxcl2, Cxcl5*) ranked among the top upregulated genes across all time points, while genes associated with resident (TLF+) macrophages (*Lyve1, Cd163*) were consistently downregulated (Fig. S3B). Gene Ontology (GO) analysis of these DEGs indicated processes related to immune cell activation at days 10 and 20, skeletal system and cartilage development at day 58, and extracellular matrix (ECM) reorganization across all time points (Fig. S3C). Extending our GO analysis of day 58 and constructing a cluster tree from the top 75 significant terms revealed that differential gene expression in the late-phase joint is enriched for pathways related to axonogenesis and neurogenesis (Fig S3D). This shows that the joint-DRG communication is an important process even after resolution of inflammation. Notably, several semaphorin genes (*Sema3a*, *Sema4a*, *Sema4d*, and *Sema5a*) were enriched within GO terms associated with axon growth in the joint on day 58. Mapping their expression across joint cell types revealed that *Sema3a* was predominantly expressed by fibroblasts, while *Sema5a* was prominent in fibroblasts, pericytes, and Schwann cells. In contrast, *Sema4a* and *Sema4d* were primarily expressed by neutrophils and other immune cells (Fig. S3E). Collectively, these findings indicate that the arthritic joint undergoes profound immune and stromal remodelling prior to sprouting of nociceptive neurons, and that multiple molecular alterations persist beyond resolution of overt inflammation. The resulting joint-DRG atlas provides a valuable resource for investigating neuroimmune interactions in the CAIA model.

### Neutrophils, not macrophages, predominate in driving arthritis and pain-like behaviour in CAIA

CAIA develops independently of T and B lymphocytes^36^, with myeloid cells playing a central role in the inflammatory process ^37^. Consistent with this, our joint-DRG transcriptomic atlas reaffirmed the importance of myeloid cells in CAIA-related processes. To delineate the differential contributions of neutrophils and monocytes/macrophages in arthritis and pain-like behaviours beyond the acute inflammatory phase, we conducted targeted depletion of each cell type and assessed mechanical and cold hypersensitivity using Von Frey filaments and the dry ice stick test, respectively. Neutrophils were depleted using recently developed depletion systems [37] that achieve more rapid and sustained depletion of neutrophils compared to the classical anti-Ly6G approach and, in contrast to anti-Gr1 depletion, selectively target neutrophils without affecting other myeloid cells, such as monocytes^38,39^. Neutrophils depletion initiated on day 3 after CAIA induction with the antibody cocktail and LPS prevented the development of arthritis as well as mechanical and cold hypersensitivity (Fig. 2A). Furthermore, when neutrophils were transiently depleted for five consecutive days starting at day 3, followed by monitoring until day 20, long-term hypersensitivity was also prevented (Fig. 2B). The five-day treatment protocol was based on experiments in naïve BALB/c mice showing that this treatment window is enough to deplete blood neutrophils and reduce DRG neutrophils by 85% (Fig. S4A, S4B). When this protocol was initiated at CAIA day 3, circulating neutrophils began to repopulate by approximately day 8, which constrained the treatment duration to five days. In contrast to day 3 administration of depletion antibodies, neutrophils depletion initiated at the peak of inflammation (from day 9) did not significantly alter joint inflammation or mechanical hypersensitivity (Fig. 2C). Similarly, administration of a murinized Ly6G depletion antibody after the resolution of overt inflammation did not alter mechanical hypersensitivity (Fig. 2D).

**Figure 2.**
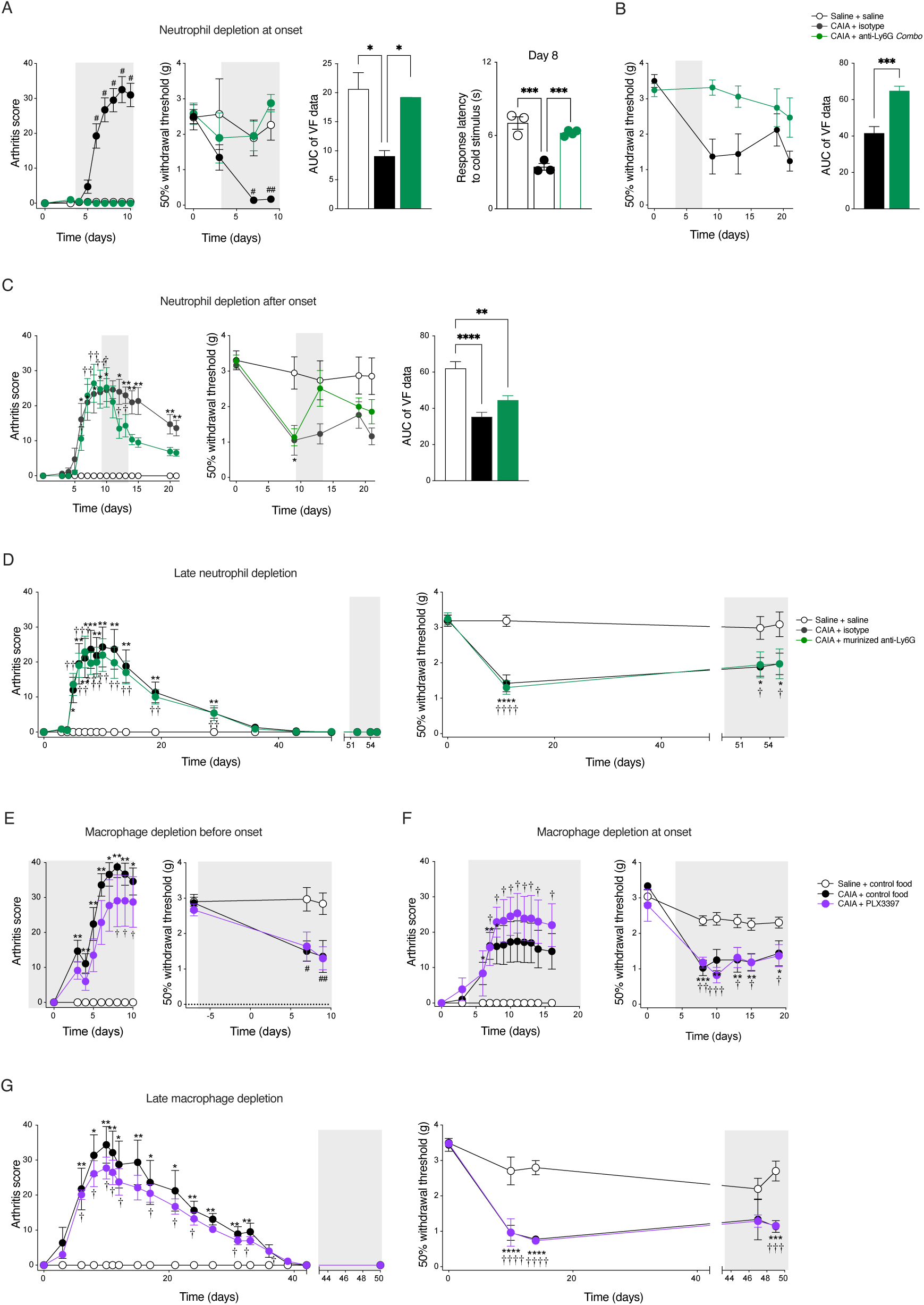
Neutrophils, not macrophages, predominate in driving arthritis and pain-like behaviour in CAIA. (A) Arthritis score, mechanical and cold hypersensitivity in saline, CAIA and CAIA mice depleted of neutrophils from day 3-10. (B) Mechanical hypersensitivity in saline, CAIA, CAIA mice depleted of neutrophils from day 3-7. (C) Arthritis score and mechanical hypersensitivity in saline, CAIA and CAIA mice depleted of neutrophils from day 9-13. (D) Arthritis score and mechanical hypersensitivity in saline, CAIA, CAIA mice depleted of neutrophils from day 51-55. (E) Arthritis score and mechanical hypersensitivity in saline, CAIA and CAIA mice depleted of macrophages. The depletion was started 7 days prior to CAIA induction and maintained through the experiment. (F) Arthritis score and mechanical hypersensitivity in saline, CAIA and CAIA mice depleted of macrophages. The depletion was started on day 4 and maintained through the experiment. (G) Arthritis score and mechanical hypersensitivity in saline, CAIA and CAIA mice depleted of macrophages. The depletion was started on day 44 and maintained through the experiment. Kruskal-Wallis test was used to compared arthritis scores. Two-way ANOVA followed by Tukeýs multiple comparison test was used for the VF data. *P<0.05, **P<0.01, ***P<0.001, ****P<0.0001. * is used for CAIA vs saline comparisons, † is used for CAIA+treatment vs saline comparisons, # is used for CAIA+ treatment vs CAIA comparisons. The grey area marks the treatment period.

The synovium contains various types of macrophages and monocytes ^13,40^, which can be targeted using the CSF1R antagonist PLX3397. We previously demonstrated that a 7-day treatment with PLX3397 was sufficient to deplete macrophages in the peripheral nervous system ^26^. In the current study, PLX3397 was provided in the mouse food at 290 ppm during three intervals: from 7 days prior to arthritis induction until day 10; from day 4 to day 21 (post-induction of inflammation); and from day 43 to day 50 (after resolution of inflammation, when pain-like behaviour persisted). Notably, depleting macrophages and monocytes did not reduce arthritis nor affect mechanical hypersensitivity, regardless of the timing of PLX3397 administration (Fig. 2E-G). Histological analysis of the DRG and joints revealed that PLX3397 treatment reduced macrophage numbers across timepoints and treatment protocols (Fig. S5A-B). Altogether, these findings identify neutrophils as key drivers of CAIA-induced inflammation and pain-like behaviour. In contrast, although macrophages undergo marked disease-associated changes in composition and organization, their depletion did not attenuate arthritis severity or pain-related behaviours, suggesting a limited pathogenic contribution to these outcomes.

### Neutrophil infiltration precedes and drives CGRP+ nerve fiber sprouting

To investigate the temporal dynamics of neutrophil-neuronal communication, we subsetted neurons and computed DEGs between CAIA and saline cells across disease stages (Fig. 3A). At day 10, neurons exhibited upregulation of gene programs associated with gliogenesis, and positive regulation of neuron projection development (Fig. 3A,B). By day 20, neuronal transcriptional changes shifted towards altered metabolic pathways and enhanced protein trafficking to the axons (Fig. 3A,B), consistent with sustained neuronal changes. In the late phase of CAIA, neurons displayed gene signatures linked to gliogenesis, glial cell differentiation and axonal ensheathment (Fig. 3A,B), suggestive of ongoing neuro-glial remodelling. Neuronal subtypes were identified using *SingleR* with a DRG atlas ^30^ as a reference (Fig. 3C). Analysis of axon sprouting–related genes across neuronal subtypes revealed increased expression in *Calca+* (CGRP) neurons compared to neuronal subtypes not expressing *Calca* (Fig. 3D), highlighting peptidergic nociceptors as a key neuronal subtype engaged during CAIA.

**Figure 3.**
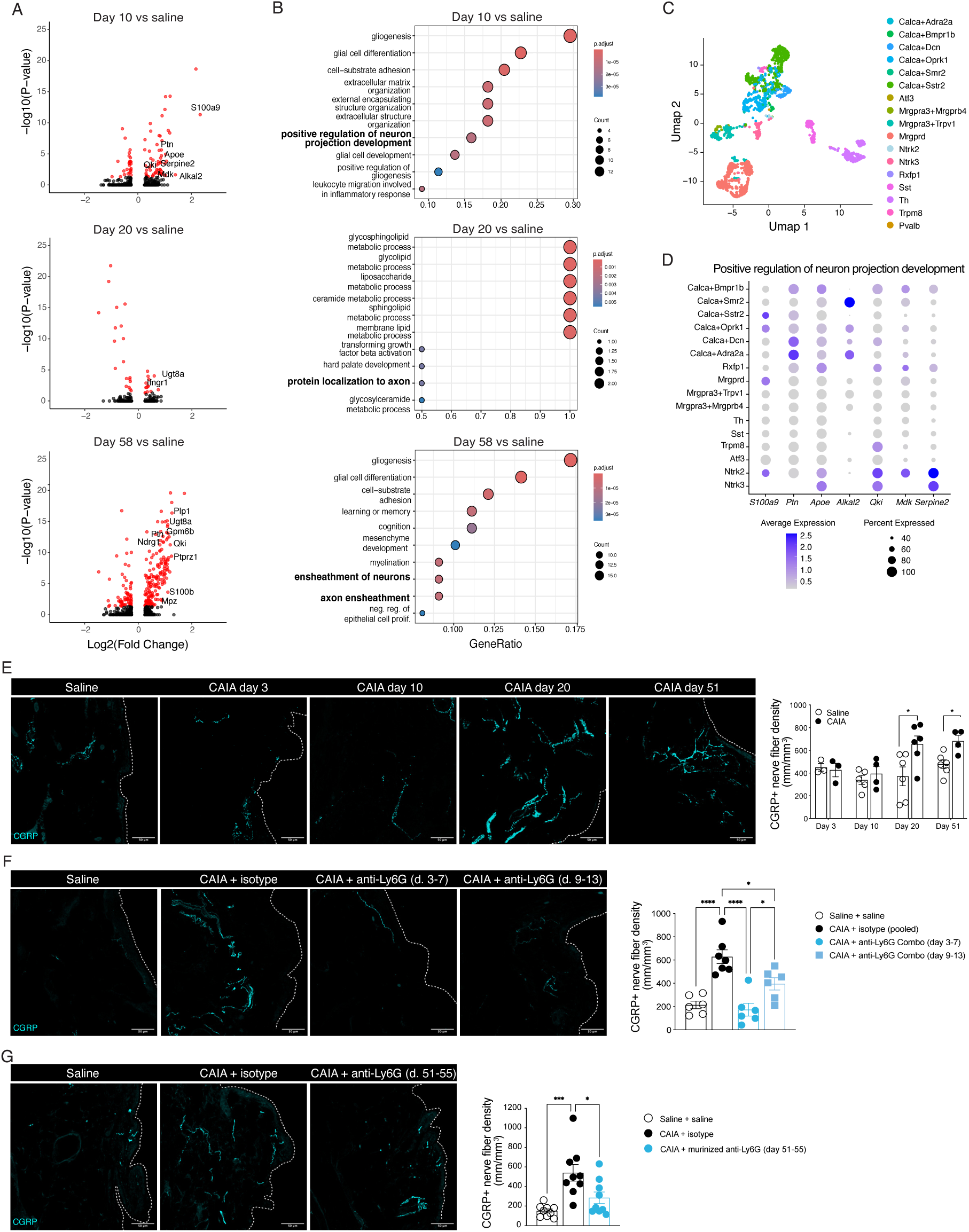
Neutrophil infiltration precedes and drives CGRP+ nerve fiber sprouting. (A) Volcano plots of differential gene expression in DRG neurons between CAIA and saline mice across timepoints. (B) GO analysis of DEGs in DRG neurons over time. (C) Subtype annotation of 2009 DRG neurons from CAIAseq based on the DRG atlas. (D) Heatmap of expression of genes listed in nerve fiber growth related terms in subtypes of neurons. (E) Stainings of CGRP+ nerve fibers in synovium of ankle joint from saline and CAIA mice at day 3, 10, 20 and 51 as well as the quantification of fibers. (F) Staining of CGRP in the ankle joint synovium of saline, CAIA and CAIA neutrophil depleted mice on day 21. (G) Staining of CGRP in the ankle joint synovium of saline, CAIA and CAIA neutrophil depleted mice on day 55. Unpaired t-test were used for saline-CAIA comparisons and one-way ANOVA was used to compare saline-CAIA-CAIA+neutrophil depletions. *P<0.05, **P<0.01, ***P<0.001, ****P<0.0001.

We previously showed that CGRP+ synovial fibers undergo robust sprouting during the late phase of CAIA, a process attenuated by janus kinase (JAK) 1/2 inhibition^6^. Given that our transcriptomic analysis revealed early activation of axon sprouting-associated gene programs in DRG neurons at day 10, we next quantified CGRP+ fiber density across disease stages using immunohistochemistry and confocal microscopy (Fig. 3E). CGRP labels multiple subtypes of nociceptive neurons that employ neuropeptides as neurotransmitters. Our data revealed that neutrophil infiltration into both the synovium and DRG preceded the emergence of synovial CGRP+ fiber sprouting, which was first detected on day 20, absent on days 3 and 10, and persisted through day 51 (Fig. 3D). Notably, transcriptomic programs associated with axonal remodelling and sprouting were already engaged at day 10, temporally coinciding with neutrophil infiltration in both tissues, suggesting that early inflammatory cues prime sensory neurons for subsequent structural remodelling.

Given that pain-like behaviour in CAIA is neutrophil-dependent and that synovial nerve fiber sprouting follows neutrophil infiltration, we hypothesized that this structural remodelling likewise requires neutrophils. To test this, we performed CGRP immunostaining on joint sections from neutrophil-depleted and isotype-treated CAIA mice, as well as from control mice. Early neutrophil depletion (days 3–7) prevented CAIA-induced sprouting of CGRP+ nerve fibers in the ankle joint synovium, resulting in normal levels of CGRP^+^ innervation in the synovium at day 20 (Fig. 3F). Although neutrophil depletion initiated at the peak of inflammation (days 9–13) did not significantly alter pain-like behaviour, it reduced CAIA-induced nerve fiber sprouting by approximately 30% reduction at day 20 (Fig. 3F). Together, these findings demonstrate that synovial nerve fiber sprouting in the CAIA model is neutrophil dependent and support a role for neutrophil-induced neuronal remodelling in the development of persistent mechanical hypersensitivity. Remarkably, depleting neutrophils after resolution of overt inflammation reduced CGRP+ fiber density by approximately 50% (Fig. 3G), indicating that inflammatory processes influencing neuronal structures remain active beyond the resolution of joint swelling.

### Neutrophils mediate sensory neuron modulation via direct and indirect mechanisms

Elucidating the molecular mechanisms underlying neutrophil-induced neuronal sprouting may enable the development of targeted pharmacological strategies that complement anti-inflammatory therapies. The increased expression of *Ifngr* in DRG neurons (Fig. 3A) raised the possibility that neutrophil-derived IFNγ could contribute to axonal remodelling, as previously reported in a nerve crush model in which neutrophil-derived IFNγ promoted nerve growth and regeneration^29^. However, *Ifng* expression was not detected in DRG neutrophils in our dataset. We therefore focused subsequent cell–cell interaction analysis on neurons and joint-infiltrating neutrophils, given the sustained neutrophil influx during CAIA. Using the R package *NicheNet*, we identified ligand–receptor pairs between neutrophils and neurons that were upregulated at day 10 of CAIA, corresponding to the onset of transcriptional programs associated with sprouting in *Calca+* neurons. This analysis revealed several candidate signalling pathways (Fig. 4A), including the *Sema4d*–*Plxnb1* axis, which has been shown to promote DRG neurite outgrowth *in vitro*^41^. We also identified oncostatin M (*Osm)* as a neutrophil-derived ligand, with its cognate receptor *Osmr* expressed by neurons. Osm-Osmr signalling engages the JAK–STAT pathway, which is of particular clinical relevance in RA given the therapeutic use of JAK inhibitors^42^.

**Figure 4.**
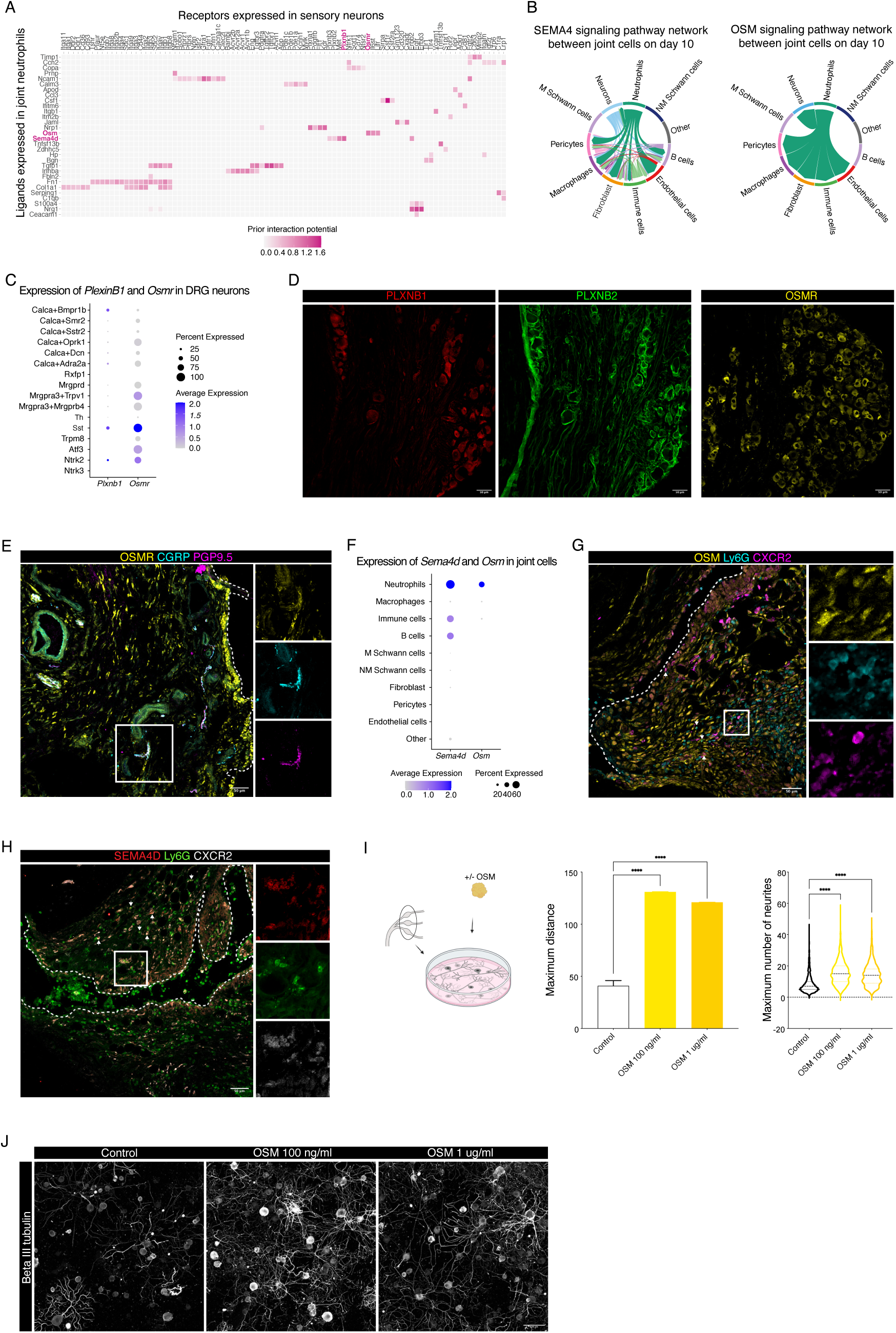
Neutrophil-neuron signalling in CAIA. (A) NicheNet analysis results of joint neutrophil to neuron signalling on day 10. (B) CellChat analysis of SEMA4 and OSM communication between joint cells and sensory neurons on day 10. (C) Expression of *PlexinB1* and *Osmr* across neuronal subtypes. (D) Stainings of PLEXINB1, PLEXINB2 and OSMR in L4 DRG neurons of naive mice. (E) Staining of healthy ankle joint synovium showing overlap in expression of OSMR with CGRP+ PGP9.5+ nerve fibers. (F) Expression of *Sema4d* and *Osm* in joint cell types. (G) Staining of OSM, Ly6G and CXCR2 in joint sections from CAIA mice day 10. Image is representative of 3 animals. Downwards arrow shows CXCR2- Ly6G^high^ neutrophils and upwards arrow shows CXCR2+ Ly6G^low^ neutrophils. (H) Staining of SEMA4D, Ly6G and CXCR2 in joint sections from CAIA mice day 10. Image is representative of 3 animals. Downwards arrow shows CXCR2- Ly6G^high^ neutrophils and upwards arrow shows CXCR2+ Ly6G^low^ neutrophils. (I-J) *In vitro* neurite growth in primary DRG cultures treated with OSM at 100 ng/ml or 1 ug/ml. Data is representing 1 experiment with 6 technical replicates from 3 animals pooled. Median values shown are calculated among all individual neurons. Between 1175 and 2135 neurons were analysed per condition. In I error bars represent the confidence interval. *P<0.05, **P<0.01, ***P<0.001, ****P<0.0001. Illustration created with BioRender.

To substantiate these findings and extend the analysis to additional joint-resident cell types, we applied the CellChat algorithm to infer intercellular communication networks. This analysis indicated that joint neutrophils engage in extensive signalling not only with neurons but also with fibroblasts, endothelial cells, pericytes, macrophages, and B cells via both Sema4d and Osm pathways (Fig. 4B and Fig. S2E), suggesting that these ligands may act as central regulators of CAIA-associated inflammatory and neuroimmune processes. Consistent with these predictions, expression analysis across neuronal subtypes revealed Plxnb1 expression in *Calca+* and *Sst+* neurons, whereas Osmr expression was highest in itch-associated neuronal populations and present at lower levels in *Calca+*, *Mrgpra3+*, and *Mrgprd+* neurons (Fig. 4C). In addition to Sema4d and Osm signalling, NicheNet analysis implicated other potential mediators of neutrophil–neuron communication, including pathways involving Ncam1, Nrg1, and Tgfb1 (Fig. 4A), further highlighting the complexity of inflammatory signalling networks influencing sensory neuron remodelling.

Immunohistochemistry confirmed protein expression of PlexinB1, PlexinB2, and OSMR in DRG neurons (Fig. 4D), supporting a functional role for these ligand-receptor pathways contribute to the neuroimmune regulation of CAIA-induced hypersensitivity. PlexinB2 expression was also prominent in satellite glial cells (Fig. 4D), indicating that SEMA4D signalling may additionally target non-neuronal components of the sensory ganglia. In the ankle joint synovium, OSMR was detected in CGRP+ sensory fibers as well as in lining and sublining fibroblasts and the vascular compartment (Fig. 4E), consistent with a broader role for OSM signalling within the inflamed joint microenvironment. Analysis of *Sema4d* and *Osm* expression across joint cell types revealed widespread *Sema4d*, with the highest levels in neutrophils, whereas *Osm* expression was more restricted and strongly enriched in neutrophils (Fig. 4F). Consistent with these transcriptomic findings, immunostaining of day 10 joint sections demonstrated OSM and SEMA4D co-localization with CXCR2+ Ly6G^low^ as well as CXCR2- Ly6G^high^ neutrophils (Fig. 4G, H).

Functionally, application of OSM to primary DRG neuronal-enriched cultures robustly promoted neurite outgrowth at both tested concentrations (Fig. 4I-J). SEMA4D has previously been reported to promote axonal growth in embryonic DRG cultures^41^.

### SEMA4D and OSM are conserved translational mediators in human RA synovium

To assess the translational relevance of the identified neutrophil-neuron signalling pathways, we analysed two independent publicly available scRNA-seq datasets: one profiling synovial cells from inflammatory arthritis patients^18^ and a second comprising a comprehensive postmortem DRG neuronal atlas^12^. The human synovial dataset exhibits a cellular composition broadly comparable to that observed in CAIA, with the expected presence of chondrocytes and a small population of adipocytes, alongside a markedly expanded lymphoid compartment including T cells and NK cells (Fig. 5A). Consistent with our murine observations, *SEMA4D* and *OSM* were expressed across multiple synovial cell types but were most highly enriched in neutrophils (Fig 5A). This expression pattern was verified by immunohistochemistry of a synovial biopsy section from the knee joint of a RA patient (Fig. 5B).

**Figure 5.**
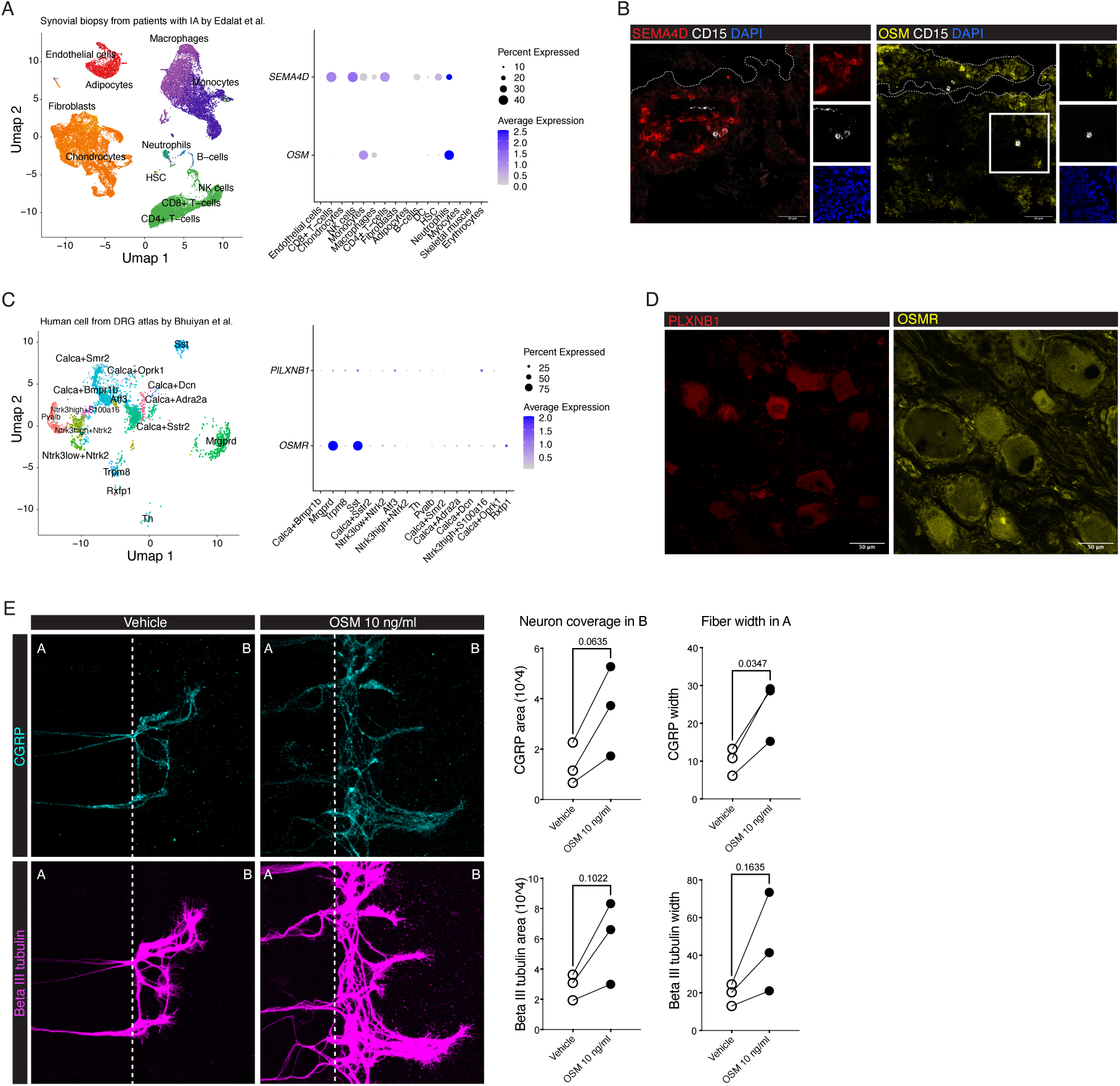
Human neutrophils and sensory neurons express SEMA4D and OSM and corresponding receptors. (A) Dimplot of synovial cell dataset published by Edalat et al. Expression of SEMA4D and OSM in human inflammatory arthritis synovial neutrophils. (B) Stainings of SEMA4D and OSM in CD15+ neutrophils in synovial sections from a knee joint biopsy of a RA patient. (C) Dimplot of human sensory neurons from the DRG atlas by Bhuiyan et al. Expression of PLEXINB1 and OSMR in human DRG neurons. (D) Stainings of PLEXINB1 and OSMR in human DRG neurons. (E) Mature HD10.6 cells treated with OSM 10 ng/ml or vehicle for 24 hours and stained for CGRP and βIII-tubulin. Individual dots represent individual experiments which are average values of three technical replicates. Paired t-tests were used for comparisons.

As scRNA-seq data from DRG tissues of RA patients are not publicly available at this time, we utilized the recently published multispecies DRG atlas^30^, restricting the analysis to human cells and nuclei (Fig. 5C). Expression mapping of the receptors *PLEXINB1* and *OSMR* showed patterns closely mirroring those observed in mouse DRG, *PLEXINB1* is expressed at low levels across several neuronal subtypes, including CGRP-expressing *CALCA+BMPR1B+* and *CALCA+SSTR2+* neurons, whereas OSMR showed higher expression in *MRGPRD+* and *SST+* neurons, with lower-level expression in *CALCA+* populations (Fig. 5C). These conserved receptor expression patterns were further validated at the protein level by immunohistochemistry on human DRG sections from organ donors (Fig. 5D).

To functionally assess whether OSM directly promotes neurite remodelling in human sensory neurons, we used the human-derived immortalized DRG neuron cell line HD10.6^43^. Differentiated HD10.6 neurons cultured in microfluidic chambers were exposed to 10 ng/ml OSM for 24 hours. Immunostaining for CGRP and βIII-tubulin revealed enhanced neurite growth in OSM-treated cultures, characterized by increased CGRP-positive fiber area in the distal compartments and greater fiber thickness within the interconnecting microchannels (Fig. 5E). Together, these findings demonstrate that OSM is sufficient to induce neurite outgrowth in human DRG-derived neurons, supporting the concept that OSM-driven neuronal remodelling is conserved across species and may represent a translatable therapeutic target for RA-associated pain.

## Discussion

Persistent joint pain remains a major unmet clinical need in RA, with a substantial proportion of patients reporting significant pain despite effective suppression of inflammation by DMARDs^1^. This dissociation between inflammatory burden and pain severity has driven increasing recognition of residual pain states in RA, implicating mechanisms beyond synovitis alone. Using the CAIA model, which isolates the effector phase of this disease^37^, we have previously shown that central sensitization and neuroimmune interplay play a role in sustaining long-lasting hypersensitivity^2,44,45^. Here we demonstrate that neutrophils are essential not only for joint inflammation but also for the induction and long-term maintenance of pain-like behaviour and pathological sensory nerve remodelling. Although neutrophils are known contributors to inflammation in CAIA ^37^, their specific role in pain-like behaviours and joint hyperinnervation has not been defined. Using a depletion strategy with improved specificity for neutrophils^38^, we show that neutrophils are required for the induction of arthritis, mechanical and cold hypersensitivity, and pathological CGRP+ synovial hyperinnervation. Notably, transient depletion early in disease was sufficient to prevent the development of long-lasting hypersensitivity, consistent with a model in which early neutrophil-dependent signals initiate durable changes in peripheral sensory circuits.

In contrast, depletion of macrophages using a CSF1R antagonist across multiple windows, before, during inflammation and after resolution, did not alter arthritis severity or mechanical hypersensitivity. This finding does not exclude key roles for macrophages in other pain contexts ^46,9^, but underscores that the contribution of myeloid subsets to nociception is highly model- and niche-dependent. Indeed, lining macrophage subsets can exert protective barrier function in K/BxN induced arthritis^47^, whereas other models implicate CX3CR1+ lining macrophages as early synovial activators that promote neutrophil recruitment^48^. In CAIA, we observed persistent disruption of the macrophage barrier niche, including reduced CD163+ sublining macrophages. This parallels evidence that CD163 deficiency worsens CAIA inflammation^49^ and clinical associations linking altered ratios of resident versus blood derived macrophage in synovium to relapse risk in RA^19,50^. Together with the limited efficacy of CSF1R and CCR2-targeting approaches in RA phase II studies^51,52^, these observations support the idea that therapeutic strategies may need to prioritize neutrophil-derived pathogenic programs while preserving or potentially restoring homeostatic CD163+ synovial macrophage functions.

Beyond immune cells, the joint stromal compartment also undergoes profound and persistent remodelling in CAIA. The pronounced shifts in synovial fibroblast populations likely reflect disruption of the synovial lining and pannus formation, hallmarks of inflammatory arthritis. Synovial fibroblasts have recently been implicated in RA pathology and in promoting synovial nerve fiber sprouting based on patient biopsies^53^, highlighting aberrant innervation as a shared feature of human RA and experimental arthritis. Nerve fiber sprouting is, thus, a common phenotype between human RA patients and arthritis models. In animal models, such nerve fiber sprouting is associated with heightened hypersensitivity and pain^53,54^. We previously reported CGRP+ synovial hyperinnervation in the late phase of CAIA^55^. Here, we demonstrate that sprouting occurs during the resolution phase of joint inflammation, follows neutrophil infiltration, and is neutrophil dependent. While sensory neurons can directly respond to immune complexes via Fc receptors^56^, our data indicate that neutrophils act as critical intermediaries linking antibody-driven inflammation to sustained neuronal remodelling and pain.

Our atlas of the joint–DRG axis in CAIA further reveals that the post-inflammatory joint fails to fully restore a homeostatic fibroblast-immune architecture, instead remaining enriched for immune-derived and extracellular matrix remodelling programs. This mirrors observations from RA patients in clinical remission, whose synovia retain molecular features associated with relapse risk^57,58^. Within this altered microenvironment, we identify neutrophil-derived OSM and SEMA4D as candidate mediators of neuroimmune crosstalk. OSM emerged as a particularly compelling factor: it is selectively enriched in neutrophils, signals via neuronally expressed OSMR, and activates JAK–STAT pathways already implicated in RA pain and therapeutically targeted by JAK inhibitors^42,59,60^. Beyond inducing neurite outgrowth in mouse and human DRG-derived neurons, OSM has previously been shown to directly sensitize sensory neurons and increase neuronal excitability^61,62^, suggesting that it may couple structural and functional plasticity to sustain hypersensitivity. These findings provide mechanistic insight into why JAK inhibition alleviates pain and reduces pathological innervation in late CAIA^6^, and they position OSM–OSMR signalling as a plausible upstream driver of these effects. Importantly, the modest efficacy of OSM-targeting antibodies in RA clinical trials^63^ may reflect trial designs focused primarily on inflammatory endpoints rather than pain or neural remodelling, rather than a lack of biological relevance. Our data therefore argue that OSM-driven neuroimmune mechanisms represent a distinct and potentially tractable dimension of RA pathology.

SEMA4D has been shown to exert anti-arthritic effects in experimental arthritis models^64^ and to promote neurite outgrowth in embryonic DRG neurons^41^, raising the possibility that it may contribute to synovial nerve remodelling. Similar to the OSM–OSMR axis, SEMA4D is expressed by synovial neutrophils from RA patients, and its receptor PLXNB1 is present on human DRG neurons, supporting translational relevance. Beyond neurons, SEMA4D has been implicated in regulation of blood–brain barrier permeability^65^, and our *in silico* analysis predict signalling to pericytes and fibroblasts, suggesting that SEMA4D may coordinate multi-compartment remodelling in CAIA. Notably, receptor expression patterns indicate pathway specificity across sensory neuron subtypes: *PlxnB1* is expressed in *Calca*+ neurons, while *Osmr* is more prominent in nonpeptidergic DRG neurons, with both receptors detectable in *Sst*+ neurons implicated in itch-related programs^12^. This segregation implies that SEMA4D and OSM may act on distinct subsets of joint-innervating sensory neurons, highlighting the importance of further investigating the neuron subtype-specific contributions to neuroimmune remodelling and pain during CAIA.

Recent studies have expanded the role of sensory neurons beyond pain transmission, implicating them in regulation of tissue immune responses in lymph nodes^66^, wound healing^67^, and cancer^68,69^. These findings raise the possibility that synovial nerve sprouting represents an adaptive neuronal response to persistent immune activation, which may in turn promote long-term hypersensitivity, particularly given that the immune landscape of the joint remains altered even at late stages of CAIA. However, this interpretation is tempered by evidence that joint inflammation is following sciatic nerve transection, suggesting that neurogenic signals are required for initiating inflammation^70^. Moreover, in CAIA mice lacking CGRP, inflammation is reduced, despite prolonged bone erosion ^71^. Together, these studies underscore a context-dependent and complex neuroimmune interplay that likely shapes both the initiation and chronicity of arthritis.

While *Osm* is preferentially expressed by joint-infiltrating neutrophils and is therefore likely to act predominantly at peripheral nerve terminals, *Sema4d* is expressed by both synovial and DRG/meningeal neutrophils, suggesting signalling at multiple anatomical levels, including distal terminals and neuronal cell bodies within the DRG. The observation that neutrophil depletion initiated after the onset of inflammation remains effective in reducing synovial nerve fiber density further supports a sustained contribution of neutrophils to nerve remodelling. Interestingly, the DRG and spinal meninges harbour a diverse neutrophil compartment even in saline-treated mice, consistent with reports showing that these cells originate from vertebra bone marrow via direct channels to the spinal cord and dura^72^. Meningeal neutrophil can be stratified into subclusters characterized by high expression of *Cxcr2*, *Cd177*, or *Orm1*, all of which expand during CAIA. Notably, Orm1+ DRG/meningeal neutrophils co-express immunoregulatory molecules such as *Cd55* and *Pgtr1*. CD55+ neutrophils have recently been shown to release vesicles that promote anti-inflammatory complement regulation^73^, whereas PGTR1 limits the availability of prostaglandins and leukotrine B4^74^. The presence of this potentially regulatory neutrophil subset may therefore constrain excessive neutrophil expansion at this site, despite high collagen II expression in the meninges^75^. Finally, the functional role of meningeal neutrophils in arthritis warrants furthers investigation. Notably, CD177+ meningeal neutrophils have been shown to modulate behavioural responses in mice^68^, raising the possibility that dural neutrophils contribute to the neuropsychological and pain-related manifestations observed in RA patients^76,77^. An aspect of neutrophil–neuronal communication in CAIA that remains to be explored in future studies is the effect of neutrophil extracellular traps-derived mediators on neuronal sprouting and excitability.

A limitation of our study is the exclusive use of female mice. This choice was driven by practical considerations, as male BALB/c mice in our facility exhibit a low incidence of robust CAIA induction, and their frequent aggressive behaviour introduces variability that can confound experimental outcomes. Although context-dependent and subject to ongoing debate, several studies have reported sex-specific differences in immune cell contributions to pain, including macrophage-dependent mechanisms ^78^. Thus, we cannot exclude that macrophages play a more prominent role in pain modulation in male mice. Nevertheless, given the higher prevalence of RA in women, the use of female animals provides clinically relevant insights into RA-associated pain mechanisms.

Another limitation is the use of a single arthritis model, which primarily captures the effector phase of arthritis through lymphoid-independent mechanisms and therefore does not fully recapitulate the adaptive immune signalling that characterizes human RA. Nevertheless, treatment-naïve RA is a heterogeneous disease, with synovial cellular architecture classified into lympho-myeloid, diffuse myeloid, and pauci-immune pathotypes, each associated with distinct treatment responses^79^. This heterogeneity highlights the importance of understanding myeloid-driven mechanisms in arthritis, while also emphasizing that different experimental models may be required to interrogate distinct disease phenotypes.

In summary, our study identifies neutrophils as central drivers of pain-like behaviour and pathological sensory nerve remodelling in autoantibody-induced arthritis, acting independently of macrophage depletion in female mice. By integrating a temporally resolved single-cell atlas of the joint–DRG axis with functional perturbation and cross-species validation, we uncover OSM and SEMA4D as conserved neutrophil-derived mediators that promote aberrant neuronal remodelling. These findings support a model in which early innate immune signals imprint durable changes in joint-innervating sensory circuits that persist beyond overt inflammation, providing a mechanistic framework for residual pain in rheumatoid arthritis. More broadly, our work highlights neutrophil-neuron communication in the synovial niche as a tractable therapeutic axis and suggests that targeting inflammation-induced neuronal remodelling in the joint may complement anti-inflammatory strategies to more effectively treat chronic arthritis-associated pain.

## Methods

### CAIA induction

Collagen antibody-induced arthritis (CAIA) was initiated by injecting 1.5 mg (150 ul) anti-CII antibody cocktail (Chonrex Inc) intravenously followed by an intraperitoneal injection of LPS from E. coli LPS serotype B4:0111 (Chondrex Inc) 20 ug in 100 ul 3 days later. In studies with significant inflammation detected already day 3, the LPS was dosed based on the arthritis score as follows: mice with a score of 0 received 25 ug, 1-9 received 20 ug LPS, 10+ received 10 ug. The mice in experiments with pre-treatment or treatment from day 3 received the same of LPS regardless of arthritis score in order to not skew the groups. Respective controls were injected with 150 ul saline intravenously and 100 ul saline intraperitoneally. From a pain perspective, the CAIA model displays two distinct phases: an inflammatory phase marked by mechanical and cold hypersensitivity, followed by a later phase where joint edema is no longer visible, yet mechanical hypersensitivity persists ^2,6,80^. This transition typically occurs 3-4 weeks after induction.

Mice that reached humane endpoints (>15% weight loss or arthritis scores >51) before the planned endpoint were excluded. Non-induction was classified as a mean arthritis score below 5 with no pain behaviour during flare unless otherwise noted. In studies with pharmacological treatments induced before expected visible inflammation, all animals were included regardless of induction (pre-treatment or treatments starting day 3-6). For experiments assessing only arthritis scores, animals with inflammation ≤2 were excluded.

### Arthritis score

Joint inflammation was assessed by scoring the inflamed joints on front and hind paws. The ankle joint and paws were scored 0, 2.5 or 5 and each inflamed digit was scored 1. The data shown represents the sum of scores on all 4 legs.

### Macrophage depletion

PLX3397 0.29 g/kg (CSF1R antagonist, Safe Augy) was introduced to mice via the food pellets. Control mice were given matching food (Safe Augy) without the drug. Treatment length was as indicated with gray boxes in individual graphs and figure legends.

### Neutrophil depletion

Neutrophils were depleted by the “double antibody-based depletion strategy” published by Boivin et al., 2020[33]. Briefly, 25 ug/mouse of InVivoPlus anti-mouse Ly6G (BioXCell) was injected subcutaneously, followed by 50 ug/mouse of InVivoMAb anti-rat Kappa Immunoglobulin Light Chain (BioXCell) injected subcutaneously as well. Antibodies were diluted in their respective InVivoPure Dilution Buffers (BioXCell). Control treatment was 25 ug/mouse of InVivoPlus rat IgG2a isotype control, anti-trinitrophenol (BioXCell), followed by a dilution buffer injection. Alternatively, recombinant mouse anti-Ly6G antibodies engineered with the same properties were also used for late phase depletion [34]. The murinized anti-Ly6G (Absolute antibody) or isotype (Absolute antibody) were administered at 25 ug per mouse 3 times over 5 days.

### Mechanical hypersensitivity

Behavioural testing was done in designated rooms and mice were habituated to the testing environment prior to testing. The tester was blinded during the experiment. In experiments where behaviour was assessed, experimental groups were assigned based on baseline measurements. Mechanical hypersensitivity was assessed using the Von Frey filaments (Marstock OptiHair) and the up-down method by Chaplan et al. 1994^2^. Filaments in the range of 0.5 to 32 mN were applied to the hind paw and held for 2s. Paw withdrawal was considered a response.

### Cold hypersensitivity

Cold allodynia tests were performed by placing mice on a 4-mm-thick glass surface in individual compartments, allowing them to habituate for 3 hours to the experimental setup. A 2-ml syringe holding a dry ice pellet was used to apply the cold stimulus. The dry ice pellet was held against the underside of the glass surface, directly beneath one hind paw, and the latency to paw withdrawal was recorded, up to a maximum of 20 seconds. There was an interval of at least 8 minutes between successive stimuli applications to the same mouse, alternating between paws. Each paw was tested three times. The six values recorded per mouse were averaged.

### Preparation of single cell suspensions

Mice were euthanized via an intraperitoneal overdose of pentobarbital (338327; APL) and then perfused with ice-cold PBS. Dorsal root ganglia (DRGs) were dissected and held in cold PBS. Bilateral L3–L5 DRGs were gathered for 10X single-cell sequencing, while all lumbar and cervical DRGs were pooled for flow cytometry and FACS analysis. The tissue was digested at 37°C for 40 minutes with an enzymatic mixture containing 2 mg/ml Collagenase I (17100-017; Gibco), 5 mg/ml Dispase II (D4693; Sigma-Aldrich), and 0.5 mg/ml DNAse I (11284932001; Roche). Samples designated for scRNA sequencing were treated with 54 µg/ml actinomycin D (000025-1890; Sigma) to prevent transcriptional alterations. The resulting cell suspension was passed through a 40-µm strainer, and enzymatic activity was halted by adding 2 mM EDTA. Myelin debris was removed using a 38% Percoll gradient (GE17-0891-02; Sigma-Aldrich), followed by washing and resuspension in PBS.

### scRNA-seq

Single-cell suspensions of dorsal root ganglion (DRG) cells from 4 female BALB/cAnNRj mice per group were first treated with an FcR blocking solution (No. 130-092-575; Miltenyi Biotec) to avoid nonspecific binding. They were then stained with Calcein Red-AM (425205; BioLegend) and the LIVE/DEAD™ Fixable Violet Dead Cell Stain (L34955, Invitrogen). TotalSeq anti-mouse Hashtag antibodies (BioLegend, TotalSeq B0301-4) were used to distinguish individual samples. The 4 mice were pooled intp 2-3 hashtags. Live, single cells (Calcein+, LIVE/DEAD-) were isolated using a BD FACSAria Fusion instrument at the Biomedicum Flow Cytometry Core Facility. Library preparation and 3’ GEM-Xv4 sequencing took place at the SciLifeLab sequencing facility in Solna on a NovaSeq 6000 platform, yielding 70,785 reads per cell. The resulting 10X CellRanger output files (barcodes, features, and count matrices) were then analyzed in R Studio with Seurat. Quality control excluded cells with fewer than 700 reads or with mitochondrial content exceeding 5%. Data normalization and scaling were performed using the Seurat functions NormalizeData and ScaleData. The variance-stabilizing transformation (vst) method in FindVariableFeatures was used to identify the top 2,000 highly variable features, which were then used for principal component analysis (RunPCA). Clustering was performed by computing K-nearest and shared nearest-neighbor graphs (FindNeighbors) with 26 dimensions, followed by Louvain-based graph clustering (FindClusters) at a resolution of 0.5, guided by cluster tree analysis (Clustree). Cell types were assigned based on established marker genes.

### Re-analysis of public datasets

The DRG atlas in the RDS file provided on painseq.shinyapps.io/harmonized_painseq_v1 was subsetted to contain only human cells in R studio and the expression of marker genes was visualized using *Seurat* and *Cellchat.* The human synovium dataset was downloaded from ArrayExpress: E-MTAB-11791, ebi.ac.uk/arrayexpress/ and the expression of genes was plotted using *Seurat* and Cellchat. without modifications to the dataset.

### Flow cytometry analysis

Single-cell suspensions prepared from dorsal root ganglia (DRG) of female BALB/cAnNRj mice were first treated with an FcR blocking reagent (No. 130-092-575; Miltenyi Biotec). The cells were then labeled with Calcein AM (425201; BioLegend) and/or LIVE/DEAD™ Fixable Violet Dead Cell Stain (L34955, Invitrogen). Primary antibodies are listed in Table 1. Staining took place at 4°C for 30 minutes, after which the cells were washed and resuspended in PBS. Data were analyzed using FlowJo 10 software.

**Table 1.** Antibodies used for flow cytometry and immunohistochemistry.

| Antigen | Clone | Vendor | Cat |
| --- | --- | --- | --- |
| Flow cytometry |  |  |  |
| CD11b | M1/70 | Biolegend | 101279 |
| TCRb | H57-597 | Biolegend | 109212 |
| MHC II | M5/114.15.2 | Biolegend | 107622 |
| Xcr1 | ZET | Biolegend | 148223 |
| CD45 | 30F11 | Biolegend | 103173 |
| CCR2 | 475301 | BD | 747963 |
| Ly6C | HK1.4 | Biolegend | 128033 |
| B220 | RA3-6B2 | Biolegend | B358940 |
| Ly6G | 1A8 | Biolegend | 127643 |
| CD11c | N418 | Biolegend | 117336 |
| CX3CR1 | Sa011F11 | Biolegend | 149021 |
| CD163 | TNKUPJ | Thermo | 12-1631-80 |
| CD206 | C068C2 | Biolegend | 141705 |
| CD64 | X54-5/7.1 | Biolegend | 139313 |
| Immunohistochemistry |  |  |  |
| CD64 | AF2074 |  |  |
| CCR2 | EPR20844-15 |  |  |
| Ly6G | 1A8 | BioLegend | 127601 |
| CD68 | MCA1957 | AbDSerotec |  |
| VSIG4 |  |  | 230721 |
| PLEXINB1 |  | ThermoFischer | 23795 |
| OSMR |  | Abcam | ab232684 |
| CD15 | Carb-3 | Agilent | GA06261-2 |
| CXCR2 | Polyclonal | GeneTex | GTX14935 |
| OSM (human) | 17001 | Bio-Tehne(R&D) | MAB295-100 |
| OSM(mouse) | Polyclonal | Bio-Tehne(R&D) | AF-495 |

### Immunohistochemistry

Mice were anathematized with isoflurane or pentobarbital and perfused cardially with PBS and 4% PFA. L3-L5 DRGs and hind leg ankle joints from hind legs were collected and postfixed in 4% PFA for 4-24 hours. Prior to blocking in OCT, DRGs were cryoprotected in 20% sucrose and sectioned at CryoStar NX70 cryostat (Thermofischer) at 10-25 um thickness. Ankle joints were decalcified in 10% EDTA and cryoprotected in 30% sucrose. Following this, they were sectioned longitudinally at 20 um. Prior to staning slides were dried 30 min at room temperature and unspecific binding was blocked using goat or donkey serum (depending on the species of the secondary antibody). Antibodies and incubation conditions used are given in Table 1. Nuclei were stained with 1:10 000 DAPI (Thermoscientific) solution for 10 min at RT and slides wee mounted with ProLong Gold Antifade Mountant (Invitrogen). Images were acquired using LSM800 laser-scanning confocal microscope (Carl Zeiss) with ZEN2012 software (Zeiss). Images were analysed using DRGquant as previously described^81^ for joint macrophage quantifications and DRG macrophages were analyzed from Z-stack maximum-projection images using ImageJ (Threshold, Fill Holes, Analyze Particles) with consistent settings for all samples. Tissue regions were defined by freehand selection on the DAPI channel. Nerve fibers were quantified using fiji by two blinded researchers.

Human lumbar DRG were collected in Canada by McGill Scoliosis and Spine team following consent from the next of kin from 4 different donors and fixed upon tissue collection for 4-6 hours or immediately frozen and post fixed for 3 hours in 4% paraformaldehyde. Tissue sections were thawed at room temperature for 1 hour, then washed twice with 0.1M phosphate-buffered saline (PBS; pH 7.35–7.45 for all PBS solutions) and once with 0.01M PBS for 5 minutes each. Next, sections were incubated at room temperature with a permeabilization solution containing 0.01M PBS, 0.16% Triton X-100, 20% dimethyl sulfoxide (DMSO), and 23 g/L glycine for 30 minutes. Following permeabilization, a blocking solution consisting of 0.01M PBS, 0.2% Triton X-100, 6% normal donkey serum, and 10% DMSO was applied for 1 hour, followed by overnight incubation of primary antibodies against OSMR (1:50, Rabbit, cat: ab232684, Abcam), PlexinB1(1:50, Rabbit, cat: 23795-1-AP, Thermofisher) at room temperature in 0.01M PBS with 0.2% Tween-20, 10 mg/L heparin, 0.02% sodium azide, 5% DMSO, and 3% normal donkey serum. After incubation, slides were washed with a washing solution (0.01M PBS, 0.2% Tween-20, 10 mg/L heparin, and 0.02% sodium azide), 1x1 min and 2x5min washes; and incubated with secondary donkey antibodies (Jackson ImmunoResearch: Cy2, 1:200; Cy3 1:600; Cy5 1:500). After secondary antibody incubation 2.5 hours, sections were washed three times using the washing solution (5 minutes each), followed by incubation with DAPI (1:10,000) for 10 minutes after which, another 5 minutes wash in washing solution was followed by a 5 minutes wash in 0.1M PBS and 2 washes in 0.01M PBS. To eliminate autofluorescence, TrueBlack Plus lipofuscin quencher (Biotium, cat: 23014) was applied 1:40 on the slides for 30 minutes and followed by 3x5 minutes washes in PBS. After drying for 5 minutes, the slides were mounted using Prolong Gold Antifade reagent (Invitrogen, cat: P36934).

Fresh-frozen synovium sections from primarily diagnosed RA and OA patients were collected from knee biopsies obtained during joint replacement surgery and cut at a thickness of 20um. The patient’s synovium sections were thawed for 15min and fixed in 4% pre-cold Formaldehyde for 20 min, then washed with 0.1M for 10 min thrice. For the OSM staining, slides were blocked with 1% BSA in 0.1M PBS for 30 min, while for the SEMA4D staining, slides were blocked in 0.1M PBS containing 0.3% Triton X-100 and 3% donkey serum for 2 hours at room temperature. Primary antibody incubation was performed overnight at room temperature. For the OSM-CD15 stainings, CD15 (Dako, GA062) and OSM (Bio-techne, MAB295, Clone#17001) were detected using AF488 donkey anti-mouse IgM (Jackson Immuno, 715-545-020) and AF647 goat anti-mouse IgG2a (Invitrogen, A21241), respectively. For the SEMA4D-CD15 stainings, CD15(Dako, GA062, Clone#Carb-3) and SEMA4D (Abcam, ab134128, Clone#EP3569) were detected using AF488 donkey anti-mouse IgM (Jackson Immuno, 715-545-020) and Cy3 donkey anti-rabbit whole IgG (Jackson Immuno, 711-165-152) respectively. Slides were then washed with 0.1M PBS for 10 min thrice and counterstained with DAPI (1:10000 in 0.1M PBS) for 5 min. After a final 0.1M PBS washing, slides were dehydrated in an ethanol gradient (70, 96, and 100% each 2 min) and rinsed twice in xylene for 2 min. Cover slipping was performed with Pertex mounting medium (Histolab). Representative images were captured in a Zeiss LSM800 laser-scanning confocal microscope with a 20x objective and a 63x objective. The region of the synovium lining and adjacent area was defined as the region of interest. The figures show an extended depth of focus projection from 21 z-stack images processed by the ZEN 3.6 software.

#### *In vitro* sprouting mouse DRG

About 40 DRGs from 3 female BALB/c mice were collected in cold HBSS. The DRGs were incubated with a pre-warmed HBSS solution containing Papain (40U/ml, Worthington), 2 ul/ml of saturated NaHCO3, L-Cysteine (0.67 mg/ml, Sigma) for 30 min at 37C. They were spun down at 300g for 2 min and incubated with a pre-warmed HBSS solution containing Collagenase Type II (4 mg/ml, Worthington) and Dispase II (4,66 mg/ml, Sigma) for 30 min at 37C. Gentle shaking was applied during both incubations. After centrifugation at 300g for 2 min, the tissues were washed with 2 ml warm media and spun the same way again. Finally, the tissues were triturate in 1 ml media (F12 with 10% HI FBS and 1% PenStrep). The cells were filtered through a 100 um strainer. Media was added to plate 150 ul on 30 wells. Non-neuronal cell depletion was done by plating the cells on non-coated flatbottom plates an incubating for 1 hour and 15 min at 37C, 5% CO2. The neuron-enriched suspension cells were transferred into a precoated (Laminin, Sigma and poly-D-Lysine, Corning) glass bottom culture plate. Control media, OSM (SinoBIologics) or SEMA4D (SinoBiologics) were added to the wells. Six well precondition were plated across two plates and incubated. The next morning the cells were washed and incubated with media of the same concentrations and incubated until total incubation was 24 hours. After that the cells were washed with DPBS and fixed with 4% PFA for 10 min. After two washes with PBS the cells were incubated overnight with 1:572 Beta II tubulin (Promega) conjugated to AF488 (Abcam). The next day the cells were washed with PBS, incubated with 1:5000 DAPI and washed with PBS and imaged.

Analysis: Confocal images were initially manually processed to eliminate large background areas by a blinded researcher. An automated analysis was later performed using equal parameters for all images using FIJI. For the processing of the neuronal bodies, beta III tubulin images were auto-thresholded using the triangle method for detection of all the neurites. Using the thresholded images, a map of the neuronal somata was estimated by eliminating thin regions. Neuronal nuclei were filtered with a low and upper threshold to separate from noise and nonneuronal nuclei respectively. Colocalization of the neuronal nuclei map and somata map was used to determine the origin centroid for Voronoi labeling of each neuron extension. Neurons were then filtered for a minimum area equal to a circle of 6 um of radius to eliminate small surrounding cells. Neuronal extension was determined once more after filtering of small objects, and the neuronal objects were isolated for Sholl analysis. Diameter was estimated by determination of max inscribed circle in the neuronal map (Max Inscribed Circles plugin). Sholl analysis was performed using SNT plugin^82^ with an initial radius of 6um and measurements every 5 um.

#### *In vitro* sprouting human DRG

Briefly, mature (D10) HD10.6 cells within microfluidic chambers (450 um microgrooves) were treated (soma & axons) for 24 hours with 10 ng/mL human OSM or vehicle control. They were then fixed and stained for DAPI, Beta III tubulin (488, green), NeuN (594), and CGRP (647, red). The end of the microgrooves and the peripheral terminal was imaged at 60X. All images were taken with the Leica SP8 confocal at the same intensities/parameters through 10 Z-stacks.

Analysis: Multi-channel fluorescence images were analyzed using FIJI (v1.54p) with the CLIJ2 package (v2.5.3.1). For true signal detection, each channel underwent Gaussian blur (σ = 1 pixel), fixed lenient intensity thresholding, and median filtering (r = 1 pixel). The three largest contiguous regions were retained and subjected to fixed strict thresholding to define positive signal areas.

Manually, blinded masks were defined to separated images into channel region and free region. For non-β-III tubulin markers, an additional β-III tubulin signal mask was applied. Quantification was performed using CLIJ2 statisticsOfLabelledPixels for both regions.

### Statistics

Unless indicated otherwise, data is shown as mean + standard error of mean (SEM). Two-way ANOVA followed by Tukeýs multiple comparison test was used for the VF data. Arthritis scores were compared using Multiple Mann-Whitney tests, followed by Two-stage step-up correction for multiple comparison (Benjamini, Krieger, and Yekutieli) when comparing two experimental groups. Kruskal-Wallis test was used to compared arthritis scores in experiments with three experimental groups. One-way ANOVA followed by Tukeýs multiple comparison test was used for significance testing of the IHC data. Unpaired t-tests were used for analysis of cell counts from flow cytometry data.

## Acknowledgements

We would like to acknowledge the contributions of the Biomedicum Flow cytometry Core facility, particularly Dr. Alda Saldan, financed by the Infrastructure Board at Karolinska Institutet, for providing access to cell sorting services, technical expertise and scientific input. Further, we would like to acknowledge the McGill Scoliosis and Spine team for providing the human DRGs. Last, we would like to acknowledge the National Genomic Infrastructure Sweden for their help with library preparation and sequencing.

