## Supplemental_figures for "Neutrophil-neuronal crosstalk drives arthritis-induced pain"

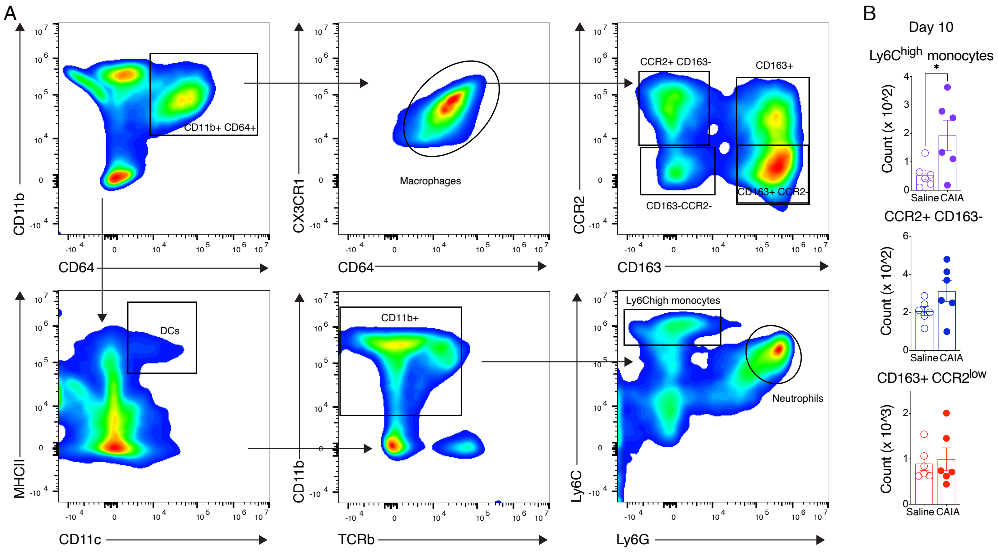


**Figure S1 Extended analysis of CAIA flow cytometry data.** (A) Gating strategy for analysis of DRG myeloid cells. (B) Counts of DRG macrophages and monocytes on CAIA day 10. Unpaired t-test were used for comparisons. *P<0.05, **P<0.01

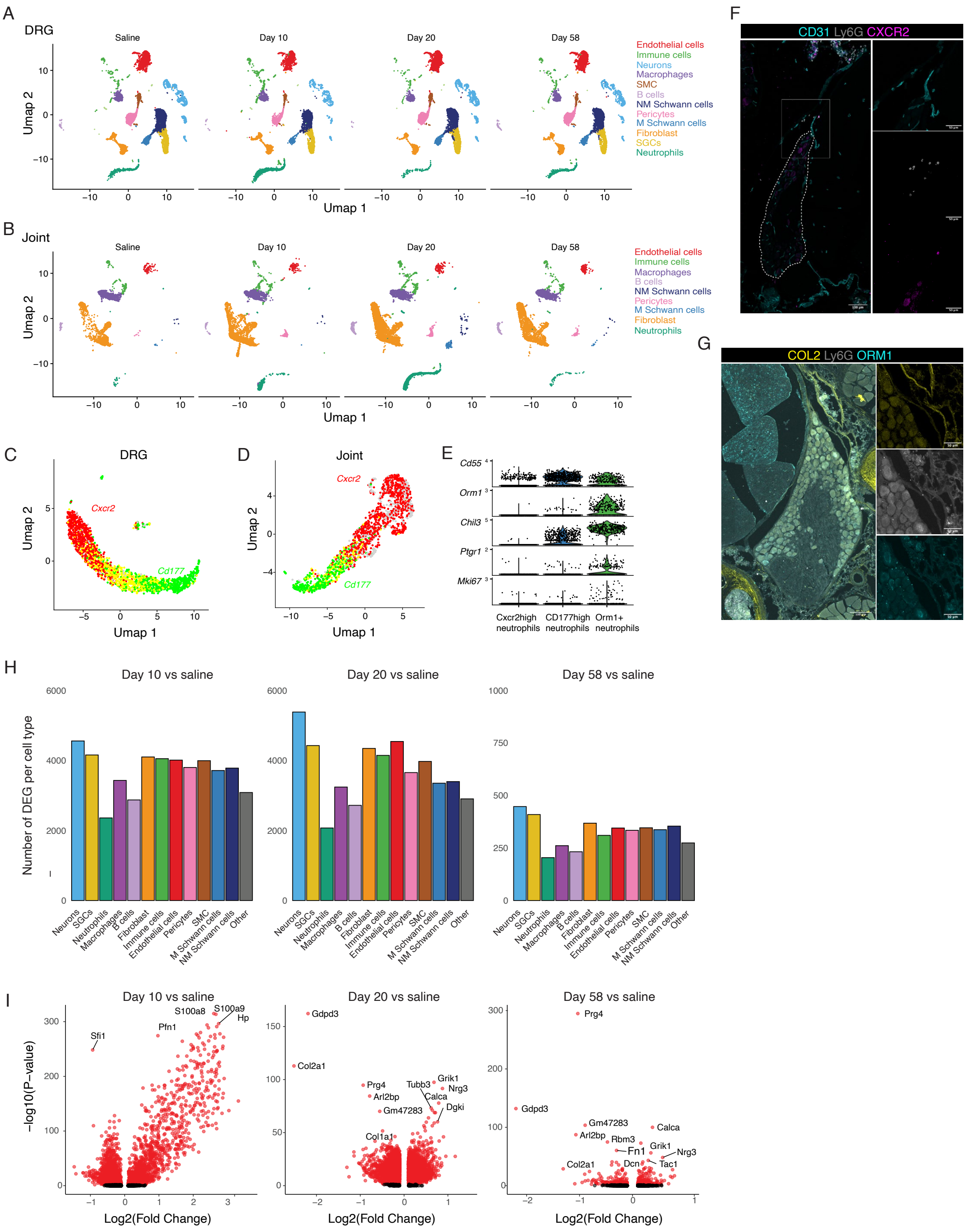

**Figure S2 Extended analysis of scRNA transcriptomic data.** (A) Umap showing the DRG cell clusters over time. (B) Umap showing the joint cell clusters over time. (C) Umap showing the expression of *Cxcr2* (red) and *Cd177* (green) in DRG neutrophils. (D) Umap showing the expression of *Cxcr2* (red) and *Cd177* (green) in joint neutrophils. (E) Expression of genes related to reduction of inflammation in DRG neutrophil subclusters. (F) Immunohistochemistry for Ly6G (neutrophils), CXCR2 (subset of neutrophils) and CD31 (blood vessels) in DRG meninges. (G) Immunohistochemistry for Ly6G (neutrophils), ORM1 (subset of neutrophils), and Collagen II DRG meninges. (H) Number of DEGs expressed in each DRG cell type. (I) Volcano plots of differential gene expression analysis in DRG cells across timepoints compared to saline. MAST was used to compute the DEGs.

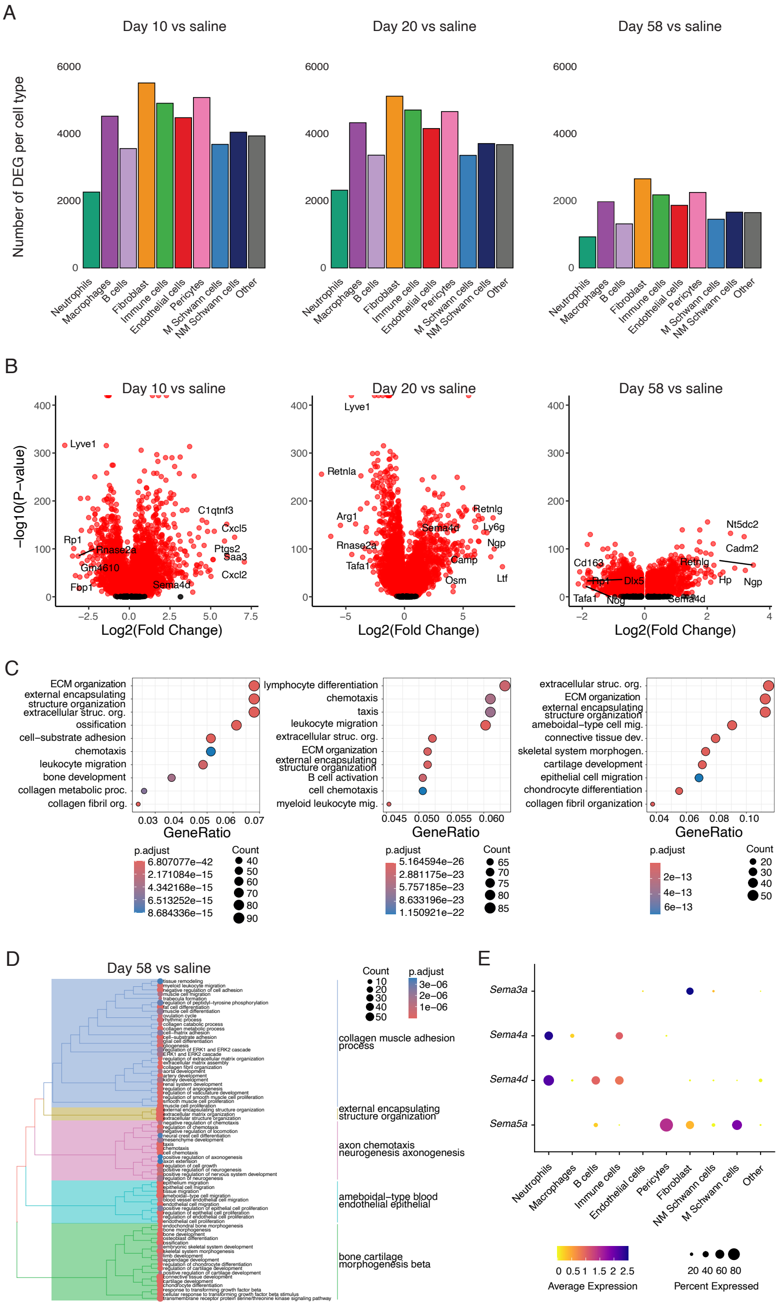


**Figure S3 Extended analysis of joint scRNA transcriptomic data.** (A) Number of DEGs expressed in each joint cell type. (B) Volcano plots of differential gene expression analysis in joint cells across timepoints compared to saline. MAST was used to compute the DEGs. (C) GO terms based in differentially expressed genes. (D) Cluster tree of 75 top terms related to DEGs between CAIA day 58 and saline cells. (E) Expression of semaphorin genes across joint cell types.

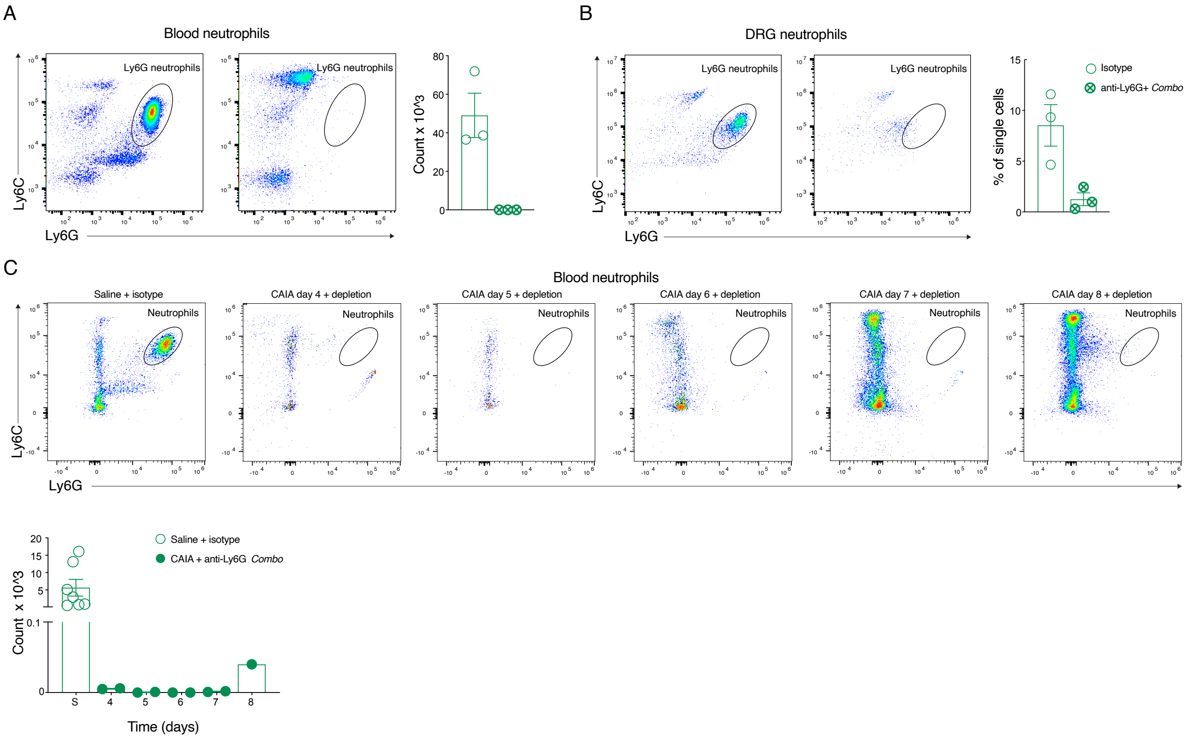


**Figure S4 Neutrophil depletion kinetics.** (A) Neutrophil counts in blood of mice treated with the *Combo* protocol for 5 days and control mice. B) Percentage of neutrophils of single DRG immune cells of mice treated with the *Combo* protocol for 5 days. (C) Neutrophil counts in blood of control mice versus CAIA mice with Combo depletion start from day 3. Time on x-axis refers to CAIA induction.

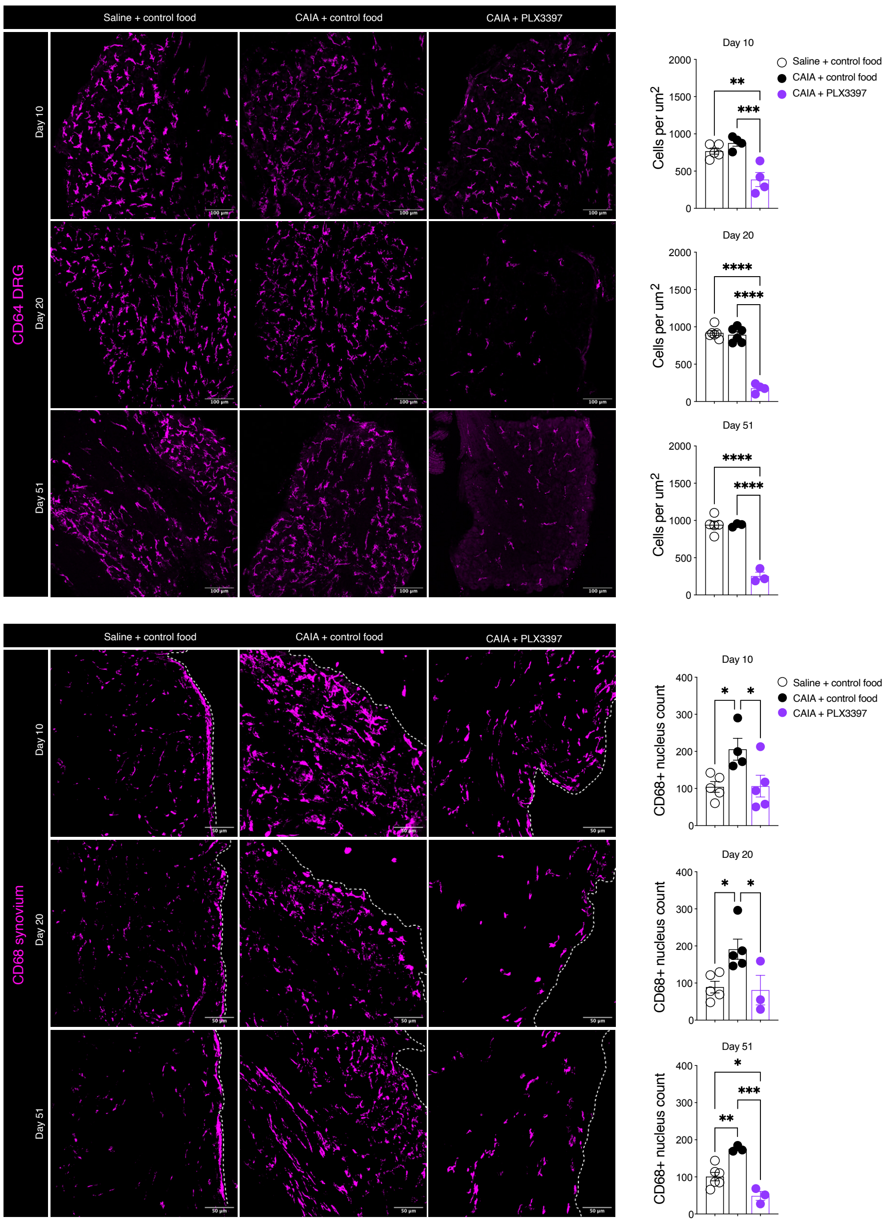


**Figure S5 Macrophage quantifications in PLX treated CAIA mice.** (A) Stainings and quantifications of CD68+ cells in the synovium of saline, CAIA and CAIA mice feed with PLX. (B) Stainings and quantifications of CD64+ cells in the DRGs, CAIA and CAIA mice fed with PLX3397. One-way ANOVA followed by Tukey´s multiple comparison test was used for comparisons. P < 0.05, **P < 0.01, ***P < 0.001.
